# Broadband high frequency Activity initializes Distractor Suppression

**DOI:** 10.1101/2024.08.22.609149

**Authors:** Paul Schmid, Christoph Reichert, Mandy V. Bartsch, Stefan Dürschmid

## Abstract

Selective attention requires fast and accurate distractor suppression. We investigated if broadband high-frequency activity (BHA; 80 – 150 Hz), indicative of local neuronal population dynamics in early sensory cortices, indexes rapid processing of distracting information. In the first experiment we tested whether BHA distinguishes targets from distracting information in a visual search paradigm using tilted gratings as targets and distractors. In the second experiment, we examined whether BHA distractor processing can be trained by statistical learning. In both experiments, BHA preceded the low-frequency target enhancement (N_T_) and distractor suppression (P_D_; 1-40 Hz) components and distinguished between targets and distractors. Only the BHA but not low-frequency component amplitude correlated with participants’ performance and was higher for lateral distractors versus lateral targets. Furthermore, BHA predicted the strength of the P_D_. These results indicate that BHA initiates stimulus discrimination via distractor suppression.

**Significance Statement:** Selective attention is the net result of successful target selection and distractor suppression, representing a key mechanism of human’s everyday life. Here, we shed light on the neural mechanisms of distractor processing by showing that the early occipital broadband high-frequency activity (BHA) is linked to successful distractor suppression. Specifically, the BHA predicted individual performance and P_D_ strength, a component associated with distractor suppression. We found the BHA and later the P_D_ increased in amplitude during implicit statistical learning indicative of a trainable distractor suppression mechanism. We demonstrate that the BHA serves as an early marker for a successful initiation of distractor suppression.

## Introduction

Fast distractor suppression is a key aspect of selective attention alongside accurate target selection (Wo stmann et al., 2019). Searching for targets amid competing distractors increases reaction times (RT) and errors (Gutteling et al., 2022; Zhao et al., 2023) indicating spatial interference (Feldmann-Wu stefeld et al., 2021), which scales with the proximity to the target (Hickey & Theeuwes, 2011; Matho t et al., 2010; Mounts, 2000). Visual search for targets is facilitated by prior knowledge about the characteristics of the targets, but also about distractors (Arita et al., 2012; Donohue et al., 2018; Mu ller et al., 1995; Olivers & Humphreys, 2003; Watson & Humphreys, 2000). Hence, success in target selection depends on fast selection and suppression of distractors (Arita et al., 2012). Selection of targets from competing distractors is addressed by a growing number of studies assessing the spatial and temporal relationships of target and distractor processing (Donohue et al., 2018; Hickey & Theeuwes, 2011). In contrast to target enhancement (Eimer, 1996; Hopf et al., 2000; Luck & Hillyard, 1994; Woodman et al., 2009), neural mechanisms of distractor suppression are less understood.

The understanding of the neural mechanisms of attentional selection of visual stimuli is mostly based on event-related potentials (ERPs), such as the N2pc (Eimer, 1996; Hopf et al., 2000; Luck & Hillyard, 1994). The N2pc is a low-frequency component (1 – 40 Hz) elicited at posterior EEG electrodes/MEG sensors between 200 and 300 ms by stimulus arrays with lateral targets and distractors in opposite hemifields. Stimulus arrays containing a lateral target while the distractor is presented on the vertical meridian and vice versa can decompose the N2pc into target enhancement (N_T_) and distractor suppression (P_D_) components (Hickey et al., 2009). Previous research (Gaspar & McDonald, 2014) suggests a greater amplitude for the N_T_ than P_D_ despite the necessity of fast and efficient suppression of distracting information. This raises the question of whether there might be another initial target-distractor distinction. Broadband high-frequency activity (BHA; 80 – 150 Hz) is a crucial analytical signal in human intracranial recordings, reflecting local neuronal population dynamics (Leonard et al., 2024; Leszczyn ski et al., 2020; Ray et al., 2008), with a rapid response modulation (< 200 ms) following stimulus onset (Bartoli et al., 2019; Gerber et al., 2017; Golan et al., 2017.; Vishne et al., 2023), preceding the occurrence of microsaccades (Yuval-Greenberg et al., 2008). Furthermore, intracranial studies on attentional selection show a lateralized BHA on attended stimuli (Szczepanski et al., 2014) occurring before the low-frequency ERPs.

Here, we used a visual search array, containing target and distractor gratings with different orientation angles, to investigate whether the BHA serves as an early indicator of distractor suppression during target discrimination. Our results suggest that the BHA initiates a trainable distractor suppression mechanism.

## Methods

### Participants

After providing their informed consent, 26 subjects participated in experiment 1, where 9 were excluded from the analysis due to excessive blink and movement artifacts, resulting in a sample size of *N* = 17 (10 female, *range*: 18 – 38 years, *M* = 26.29 years, *SD* = 4.92 years). Thirty subjects participated in experiment 2 (21 female, *range*: 19 – 39, *M* = 25.47 years, *SD* = 4.03 years). Sample sizes are comparable with previous studies investigating the N2pc and its subcomponents (Gaspelin & Luck, 2018; Hilimire et al., 2011; Marturano et al., 2020). All participants reported normal or corrected to normal vision and no history of neurological or psychiatric diseases. The recordings took place at the Department of Neurology, Otto-von-Guericke University Magdeburg and were approved by the local ethics committee (“Ethical Committee of the Otto-von-Guericke University Magdeburg”). All participants were compensated with 28€.

### Experimental Design Stimuli

The participants were presented with a visual search array composed of red, green and blue grating patterns, each consisting of three stripes as viewed through a circular aperture, displayed on a gray background (see Figure 1). Red and green gratings were alternated as targets and distractors randomly between blocks, while blue gratings served as nontargets. If the red grating was the target, the green grating was the distractor and had to be ignored and vice versa. Search arrays always consisted of 19 nontargets, one pop-out target and one pop-out distractor arranged in 7 columns with 3 gratings each, and were displayed below a fixation cross to ensure a strong N2pc response (Hilimire et al., 2011; Luck et al., 1997). Participants were asked to fixate the cross, which was located at 3.75 ° visual angle (va) above the search array. The size of each grating was 0.54 ° va, and within a column they were spaced 0.01 ° va apart from each other. Target and distractor gratings could be tilted left or right in 10 steps of 1.5 °, ranging from 1.5 ° to 15 °, such that across all trials, there was a continuous distractor-target difference ranging from 0° to 30° in 21 steps. The angle of each individual nontarget was randomly determined, ranging from 0 ° to 90° (see Figure 1). Stimulus generation and experimental control was done using Matlab R2019 (Mathworks, Natick, USA) and the Psychophysics Toolbox (Brainard, 1997). Colors were matched for isoluminance using heterochromatic flicker photometry (Lee et al., 1988).

**Figure 1.**
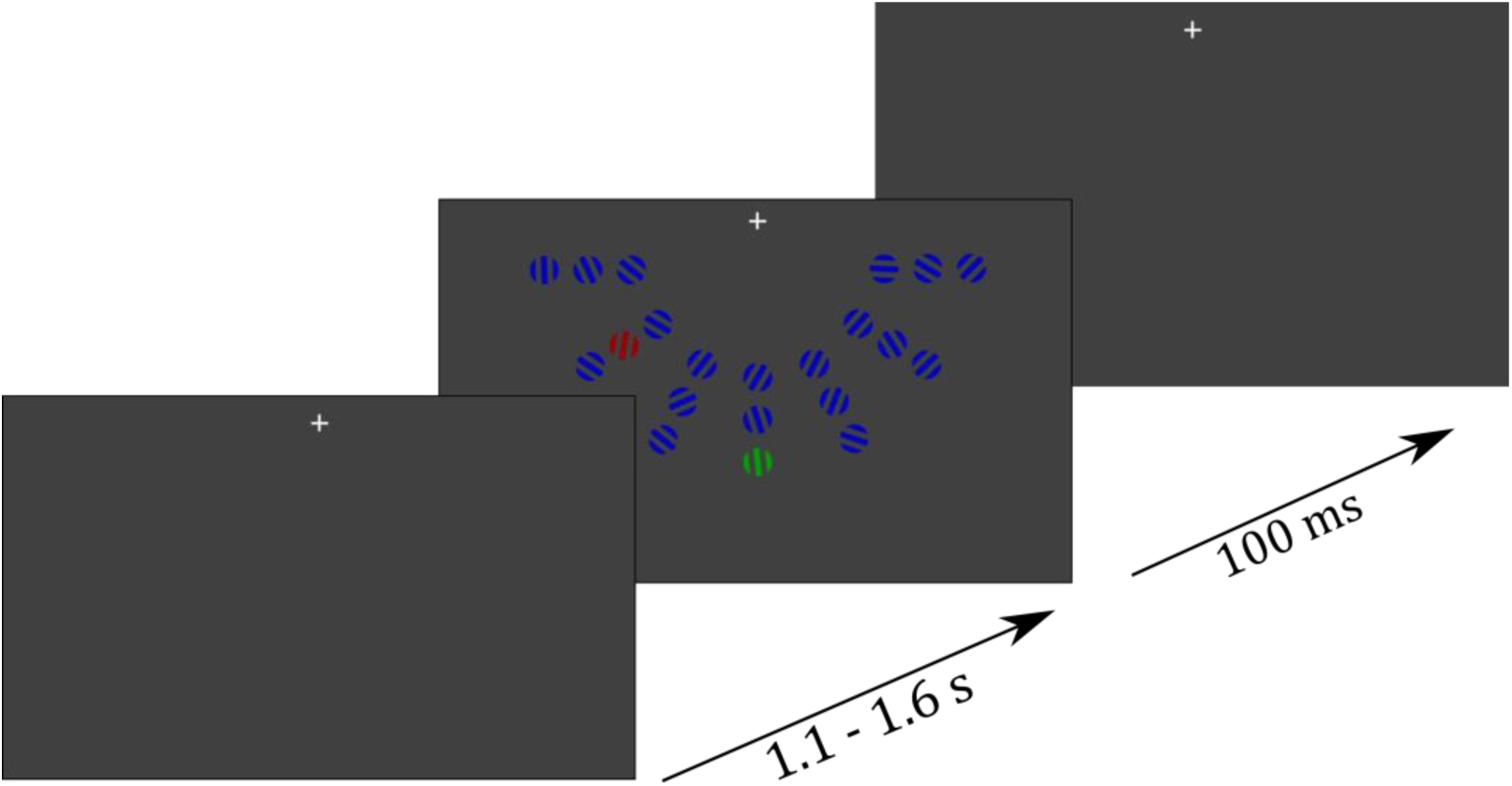
Example of a trial. Participants were presented with a fixation cross for 1.1 – 1.6 s, followed by presentation of a search array, containing 19 nontargets, 1 target and one pop-out distractor for 100 ms. In each block the target color was changed between green and red. In blocks with the green grating serving as the target the red grating served as the pop-out distractor and vice versa. After presentation of the search array, participants responded via button press whether the target grating was oriented to the left or right.

### Procedure

At the beginning of each of the six blocks, the participants were instructed to either attend the red and ignore the green grating or vice versa in both experiments. The task was to report via button press whether the target was oriented to the left or the right side using the right index and middle finger, respectively. Target color assignment varied in a pseudorandom order with 50% of the blocks containing red as the target color and the other half containing green as the target color. To distinguish between a lateral brain response to target and distractor, only one of them can be presented in a lateralized manner. This means if the target was presented on one of the three outer left or right columns, the distractor was presented on the vertical meridian. In the MEG, this leads to a lateralized target response. To determine the distractor response, the distractor was presented on one of the six lateralized columns while the target was presented on the vertical meridian column. Lateral target and distractor trials varied in a pseudorandom order with 50% of the trials containing lateral target and central distractor and vice versa. Each trial started with the presentation of a fixation cross for 1350 ms (± 250 ms) before the search array was presented for 100 ms. Participants were asked to respond as fast as possible to report the orientation of the target grating. Each of the six blocks contained 252 trials. In experiment 1 the positions of the lateralized distractors varied randomly. In experiment 2, half of the blocks functioned as implicit learning blocks in a pseudorandomized block order. Previous studies have shown that the interfering distractor effect increases with the proximity between distractor and target: reaction times slowed, discrimination performance decreased and P_D_ increased with proximity to the target (Feldmann-Wu stefeld et al., 2021; Hickey & Theeuwes, 2011). Thus, to maximize measurable distractor suppression responses, we chose the positions closest to the target for our implicit learning account. Specifically, in 65 % of trials in implicit learning blocks the distractor was presented on one of the six locations closest to the vertical meridian if the target was presented centrally (see Figure 5A). This was done to induce implicit learning of the distractor location. We assumed, that distractor suppression signals (P_D_ and BHA to lateralized distractors) for these locations should be higher in learning blocks compared to non-learning blocks if the participants implicitly learned that the distractor is likely presented close to the target.

### MEG and Eye Movement Recordings

In both experiments, participants were provided with metal-free clothing and were seated in a dimmed, magnetically shielded recording booth. Stimuli were presented on a rear-projection screen with a viewing distance of 100 cm to the participant using an LCD projector. Responses were given with the left and right hand via an MEG-compatible VPixx (VPixx Technologies, Saint-Bruno, Canada) response system. Acquisition of MEG data was performed using a whole-head Elekta Neuromag TRIUX MEG system (Elekta Oy, Helsinki, Finland), which contains 102 magnetometers and 204 planar gradiometers. Sampling rate was set to 1000 Hz. Vertical EOG was recorded bipolar with electrodes placed above and under the right eye. Horizontal EOG was recorded bipolar with electrodes placed on the left and right outer canthus. In experiment 2, we furthermore recorded participants’ eye movements in a subset of 23 participants using an Eyelink 1000 Plus system (SR Research) with a Long-Range Mount configuration and a built-in camera. We attached the Long-Range Mount configuration to the presentation screen, so that all participants were seated in the same distance to the eye tracking camera (100 cm). We recorded pupil dilation of the participants’ left eye with a sampling rate of 500 Hz. Prior to data collection, participants completed an eye tracker calibration with the built-in 9-point grid method. Preparations and recording took about 2.5 hours.

### Preprocessing

All preprocessing steps were conducted using Matlab 2013b (Mathworks, Natick, USA). Maxwell filtering was applied to reduce external noise and MEG and EOG data were down-sampled to 500 Hz. We filtered the EOG data between 1 and 40 Hz using a Butterworth bandpass filter. Trials with a variance that exceeded 4 times the mean variance were excluded. We also excluded activity stemming from eye movement artifacts from the ongoing signal, using a linear integration approach (Parra et al., 2005). Data were epoched from 1s preceding the stimulus presentation to 2 s after the presentation onset – sufficiently long to prevent edge effects during filtering. Each trial was then baseline-corrected relative to the 500 ms interval preceding the stimulus presentation. The 102 magnetometers were involved in our analyses. All filtering (see below) was performed using zero phaseshift IIR filters (fourth order; filtfilt.m in Matlab). First, we filtered the data between 1 and 200 Hz. Then, we notchfiltered the data to discard line noise (50Hz and 3 harmonics). To discard trials of excessive, nonphysiological amplitude, we used a threshold of 3pT, which the absolute MEG values must not exceed. We then visually inspected all data, excluded epochs exhibiting excessive muscle activity, as well as time intervals containing artifactual signal distortions, such as signal steps or pulses. On average, 1.6 % of the trials in experiment 1 and 1.2 % in experiment 2 were rejected due to artifacts.

### Statistical Analysis

For statistical analysis, we conducted the following analysis steps. We compared behavioral measures for lateralized targets and distractors as well as for different target orientation angles in experiment 1. Furthermore, we analyzed behavioral responses as a function of the angular difference between target and distractor. This included both reaction times and performance, where performance was defined as the percentage of correct responses (*I – Behavioral Performance*). We then tested whether the low frequency MEG N_T_ and P_D_ components varied with behavioral performance (*II – N_T_ and P_D_ Response*). Next, we identified MEG channels showing stimulus related MEG-BHA response (*III – Broadband High Frequency Activity*). We further tested whether the BHA response differed between lateral targets and distractors (*IV – BHA response to Targets and Distractors*). In a next step, we compared the BHA response characteristics to lateralized targets and distractors with their respective low-frequency counterparts (*V – Comparison between BHA Response and N_T_ / P_D_ Response*). In an exploratory analysis, we finally tested whether lateralized targets and distractors elicit a lateralized BHA response (*VI – Lateralized BHA Response*). In experiment 2, we first extracted the low frequency P_D_ and N_T_ components and extracted MEG sensors showing a stimulus-related MEG-BHA response (*VII – Stimulus Response*). We then extracted microsaccades from the eye movement data and compared the temporal evolution with the time course of P_D_, N_T_ and BHA responses (*VIII – Temporal Evolution of Microsaccades*). In a final step, we tested whether the P_D_ and BHA response differed between learning and non-learning trials (*IX – Amplitude Modulation with Experimental Condition*). To determine statistical significance, we compared each statistical parameter against a surrogate distribution. For the grand average responses of N_T_, P_D_ and BHA, we compared the observed data against a surrogate distribution as well. In 1000 iterations, we constructed the surrogate distribution by circularly shifting time series of participants between -1 and 2 s separately. We then compared the original data with this surrogate distribution by calculating the cumulative distribution function, which resulted in a *P* value for each time point indicating whether there was a significant amplitude modulation compared to the baseline.

### I – Behavioral Performance

#### Discrimination performance

We compared performance (percentage of correctly reported target orientation) between lateral and central targets. We first tested whether a greater orientation angle of the target leads to an improvement in performance. To quantify the distractor effect, we compared whether the target discrimination performance changed with the angular difference between the orientation angles of target and distractor. First, we compared performance differences for different target angles using a one-way ANOVA with the factor stimulus angle (1.5° to 15° in 10 steps of 1.5°). We then grouped performance for trials with low (≤ 7.5 °) and high (≥ 9 °) target angles and compared them using a *t*-test. To anticipate low and high target angles corresponds to subthreshold (< 70%; see Results) and suprathreshold performance (> 80%), respectively. Hence, in the following we use subthreshold and suprathreshold performance referring to low and high target angles, respectively. In a next step we tested the distractor effect in high-angle targets since only targets with high angles allow strongest angular differences (e.g., a small target angle of 1.5 ° to the left would allow only for a maximal angular difference of 16.5 ° when the distractor is tilted 15 ° to the right). Angular differences between target and distractor, ranging from 0°to 30°, were divided in two halves (low vs. high, see *Results*). We averaged participants’ performance for high target angles and compared them between low and high angular differences using a *t*-test.

#### Reaction times

We carried out the same analysis steps as for the performance comparisons. First, we compared reaction times between lateral targets and distractors using a one-way ANOVA. Post-hoc, we grouped reaction times for trials with subthreshold and suprathreshold performance target angles and compared them using a *t*-test. In a last step we averaged participants’ reaction times for suprathreshold stimuli and compared them between low and high angular differences using a *t*-test.

### II – N_T_ and P_D_ Response

The N_T_ response was determined in the following way (Boehler et al., 2011): For each participant, we averaged MEG data separately for trials with targets presented in the right and left visual field. The N_T_ was quantified in sensors showing strongest mean activity between 200 ms and 300 ms after stimulus presentation in corresponding efflux/influx zones. The signal of the sensor corresponding to the influx was subtracted from the signal of the sensors corresponding to the efflux. We then calculated the difference wave by subtracting MEG activity to targets in the right visual field form targets in the left visual field. To better compare N_T_ and P_D_ responses, we presented both P_D_ and N_T_ data as a positive going responses (by inverting the sign of the N_T_ waveform), and for better comparison of the response characteristics of the low-frequency components with the BHA, we z-standardized the low-frequency components by performing a baseline correction to correct for both zero mean and unit standard deviation in the baseline period, i.e., at each time point we subtracted the baseline mean from the low-frequency component signal and divided the result by the standard deviation of the baseline. To assess the time range of significant N_T_ response, we compared each time point of the observed modulation between 100 and 400 ms with a surrogate distribution (see *Statistical Analysis*). *P* values were then corrected for multiple comparisons, using the False Discovery Rate (FDR) method (Benjamini & Hochberg, 1995). Finally, we analyzed the relationship between participants’ N_T_ amplitude and participants’ performance using Pearson correlation coefficients, resulting in a time series of correlation coefficients. The procedure to characterize the P_D_ response followed the steps explained for the N_T_ but for trials with lateral distractors. We also analyzed the relationship between participants’ P_D_ amplitude and their performance using Pearson correlation coefficients. Finally, we compared N_T_ and P_D_ responses using a *t*-test to reveal possible differences in the neuronal response to lateralized targets and distractors in the low-frequency components.

### III – Broad Band High Frequency Activity

For each trial and channel, we band-pass filtered the time series in the broadband high frequency range (80–150 Hz) and obtained the analytic amplitude Af (t) of the signal by Hilbert-transforming the filtered time series. Afterwards we identified channels showing a significant amplitude modulation in the BHA following the stimulus presentation. Since we expected a BHA modulation within the first 300 ms, we z-standardized BHA amplitude values (see *Statistical Analysis II – N_T_ and P_D_ Response)*. Channels with z-scores higher than 2 in the interval ranging from 0 to 300 ms were labeled as stimulus responsive. We then averaged the BHA data across the significant MEG channels and determined the time window, in which the BHA showed a significant modulation by comparing each observed time point with the surrogate distribution (see *Statistical Analysis*).

### IV – BHA Response to Targets and Distractors

In a next step we tested whether the BHA distinguishes between lateral targets and distractors. We grouped BHA response for trials with lateral target and distractor. For both conditions (lateral target, lateral distractor), we determined the time interval of significant BHA amplitude modulation over baseline by comparing each observed time point with a surrogate distribution (see *Statistical Analysis*). Afterwards, we compared BHA responses to lateral targets and distractors using *t* tests. Since the BHA response is known to have a fast onset and a slowly decreasing flank, we compared mean amplitudes of the increasing BHA flank and the decreasing BHA flank separately. To investigate the relationship between the participants’ BHA amplitude and their performance, we calculated Pearsons’s correlation coefficients, resulting in a time series of correlation coefficients.

### V – Comparison between BHA Response and N_T_ / P_D_ Response

To investigate how the BHA response characteristics relate to those of the low-frequency counterparts (N_T_ and P_D_), we compared the correlation between BHA amplitude and performance with the correlation between the respective low-frequency components amplitude and performance. We further analyzed possible amplitude and latency differences between the BHA response to lateral targets and the N_T_, as well as between the BHA response to lateral distractors and the P_D_ by calculating participants’ individual time points of peak response for each component and comparing them using *t*-tests. In a final step we analyzed the direct link between participants’ mean BHA and low-frequency component amplitudes by calculating the Pearson’s correlation coefficient. We also tested whether BHA amplitude predicted P_D_ and N_T_ amplitude and calculated Pearson correlation coefficients between BHA amplitude at peak time point and P_D_ and N_T_ amplitude values at each time point separately. We compared these correlation coefficients with a surrogate distribution. In 1.000 iterations, we took the BHA and low-frequency component values of the subjects at the time of the highest observed correlation, randomly reassigned the BHA and low-frequency component values to the subjects and then calculated the Pearson correlation coefficient. We then calculated the 99% Confidence Interval for the correlation coefficient resulting in a critical coefficient value. Correlation coefficients exceeding this critical value were considered as showing a significant correlation between BHA peak response and low-frequency component amplitude.

### VI – Lateralized BHA Response

In an exploratory analysis, we investigated whether lateral targets and distractors elicit a lateralized BHA response. We first analyzed whether the grand average BHA showed a lateral response. We averaged BHA response over stimulus responsive channels (see *Statistical Analysis III – Broad Band High Frequency Activity*) located in the hemispheres contralateral and ipsilateral to the presented stimulus separately, resulting in a contralateral and ipsilateral time course of BHA response. To investigate whether BHA showed a lateral response like N_T_ and P_D_, we separated trials for lateral target and distractor, resulting in separate contralateral and ipsilateral BHA responses for lateral target and distractor. We then compared mean contralateral and ipsilateral BHA responses using *t* tests.

### VII – Behavioral Consequences of Implicit Learning

#### Discrimination Performance

We compared performance (percentage of correctly reported target orientation) in trials with lateral distractors between learning and non-learning conditions. We compared performance for all trials and for trials where the distractor was presented at the position with the highest occurrence probability. Due to the simplicity of the task, we expected a ceiling effect that could not be further improved by learning. We tested differences in the distractor effect across all target angles and all target-distractor orientation differences. We averaged participants’ performance and compared it between learning and non-learning conditions using a *t*-test.

#### Reaction Times

We carried out the same analysis steps as for the performance comparisons.

### VIII – Stimulus Response

The procedure to characterize the N_T_ and P_D_ response in experiment 2 followed the analysis steps for the N_T_ and P_D_ response in experiment 1 (see *Statistical Analysis II – N_T_ and P_D_ Response*). Analyzing the BHA response, we followed the steps to extract the BHA response in experiment 1 (see *Statistical Analysis II – Broad Band High Frequency Activity*).

### IX – Temporal Evolution of Microsaccades

In experiment 2, we used the built-in function of the eye tracker to extract saccadic eye movements. For each participant, we first epoched the ongoing signal from 1s preceding the stimulus presentation to 2 s after the presentation onset. This resulted in a matrix containing information on the temporal evolution and amplitude of saccadic eye movements for each trial. We then averaged the data across trials for each participant indicating how many saccadic events occurred on average at a given time point during the trials. Since we were interested in microsaccades, we only included saccades with an amplitude < 0.5° va. In a final step, we compared the temporal evolution of microsaccades with the low-frequency N_T_, P_D_ and BHA response. We calculated participants’ time points of peak N_T_, P_D_ and BHA amplitude as well as the time points at which the most microsaccades occurred and compared peak microsaccade time points with peak N_T_, P_D_ and BHA time points separately using *t* tests. Finally, we analyzed whether microsaccade rates differed between trials with lateral target and distractor by comparing mean microsaccade rates in the time range of the observed low-frequency components using a *t-*test.

### X – Amplitude Modulation with Experimental Condition

In experiment 2, we then analyzed whether the low-frequency P_D_ and N_T_ components and the BHA response were modulated by implicit learning. We averaged P_D_, N_T_ and BHA activity for trials with the distractor (for P_D_ and BHA) and target (for N_T_) being presented on one of the six locations closest to the vertical meridian (see Figure 5A) for learning and non-learning trials separately. We then analyzed possible differences between learning and non-learning trials in mean P_D_, N_T_ and BHA response separately using a *t-*test. Similar as in experiment 1 (see *Statistical Analysis – IV BHA Response to Targets and Distractors*), we compared mean amplitudes of the increasing BHA flank and the decreasing BHA flank separately. As we could only include a limited number of trials in the analysis of the implicit learning data due to the nature of our implicit learning approach, we z-standardized the P_D_, N_T_ and BHA time series (see *Statistical Analysis II – N_T_ and P_D_ Response)*.

## Results

### I – Behavioral Performance

#### Discrimination performance

Participants discriminated central targets better than lateral targets (*M*_central_ = 78.5 %; *M*_lateral_ = 72.7 %; *t*_16_ = 2.63; *P* = .02; see Figure 2A). A one-way ANOVA with the factor target orientation angle (1.5° to 15° in 10 steps of 1.5°) showed a significant main effect (*F*_9,160_ = 32.69; *P* < .0001) with higher orientation angle increasing performance (see Figure 2A). Performance for trials with subthreshold (≤ 7.5 °; see Methods) targets was significantly lower than for suprathreshold (≥ 9 °) targets (*M*_sub_ = 68.7 %; *M*_supra_ = 82.3 %; *t*_16_ = 19.75; *P* < .0001; see *Supplementary Table 1*). We tested whether the orientation angle difference between target and distractor modulated discrimination performance (see Figure 2A). High target orientation angles can have both a low and high angular difference relative to distractors. Hence, we evaluated target discrimination performance by examining angular differences for suprathreshold targets (≥ 9 °; see Figure 2A). We averaged performance for trials with low (< 13.5°) and high (> 16.5°) target-distractor angular difference, separately. We found a significant difference between low and high angular differences (*M*_low_ = 83.5%; *M*_high_ = 80.4%; *t*_16_ = 3.17; *P* = .006; see Figure 2A), with higher performance in low angular difference trials.

**Figure 2.**
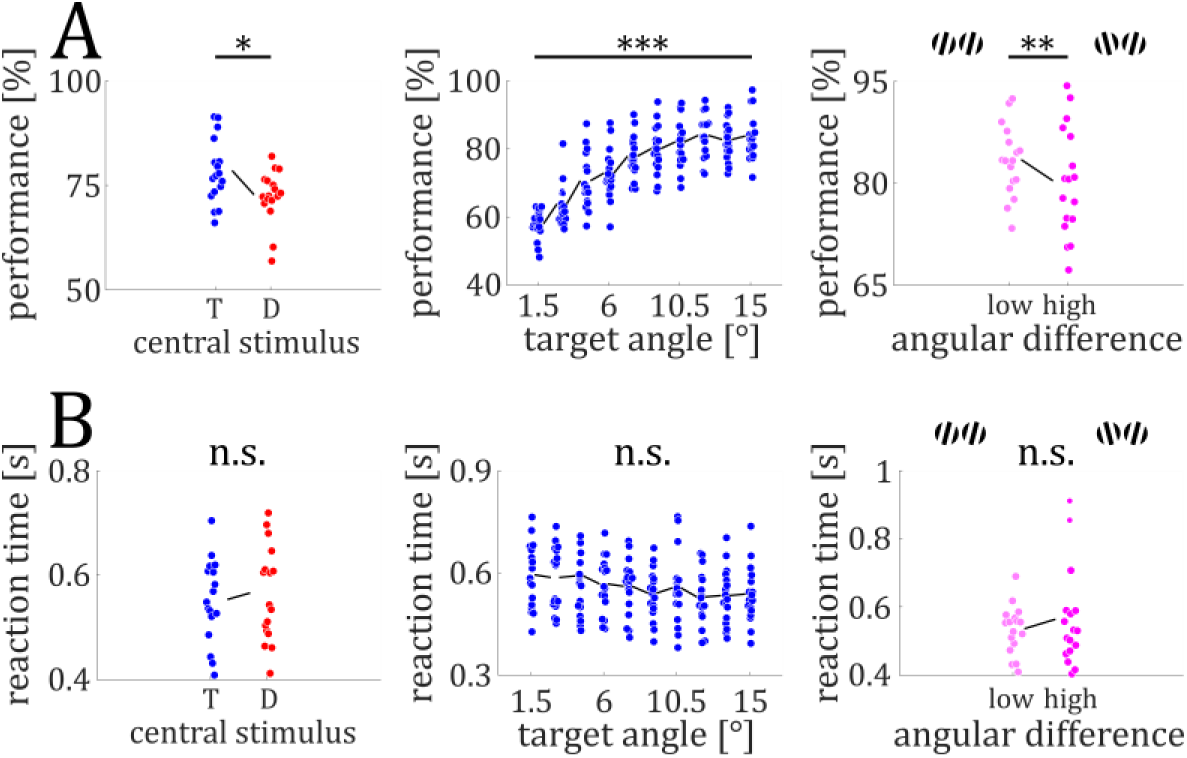
Behavioral results. ***A***: Participants discriminated central targets better than lateral targets (left). Performance increased with target orientation angle (middle). Performance for high target angle stimuli (9° to 15°) was higher for lower angular differences between target and distractor (0° to 13.5°) compared to higher angular differences (16° to 30°) (right). ***B***: No differences in reaction times between trials with central target and distractor (left). No differences in reaction times between the different target orientation angles (middle). No differences in reaction times for high target angle stimuli between lower and higher angular differences (right). Colored circles represent single data points. Error bars represent the standard error of the means (SEM). * = *P* < .05. ** = *P* < .01. *** = *P* < .0001.

#### Reaction times

Reaction times did not differ between trials with lateral targets vs. lateral distractors (*M*_Target_ = 553 ms *M*_Distractor_ = 566 ms; *t*_16_ = 1.07; *P* = .30; see Figure 2B). A one-way ANOVA with the factor target orientation angle (1.5° to 15° in 10 steps of 1.5°) did not show a significant main effect (*F*_9,160_ = 1.13; *P* = .34). In a next step, we averaged reaction times for suprathreshold target trials for low and high angular differences separately. We did not find any differences in reaction times between low and high angular differences (*M*_low_ = 536 ms; *M*_high_ = 564 ms; *t*_16_ = 1.16; *P* = .26; see Figure 2B).

### II – N_T_ and P_D_ Response

Lateral targets elicited a N_T_ response between 166 and 330 ms (z_crit_ = 3.47; N_Tmax_ = 15.24 at 258 ms; all *P* < .04; see Figure 3A). The N_T_ amplitude was not correlated with participants’ performance, neither for the time resolved analysis (*r*_max_ = .40 at 180 ms; all *P* > .11; see Figure 3A), nor for correlation with N_T_ averaged across the time interval of significant N_T_ (*r* = .24; *P* = .36; see Figure 3A). Lateral distractors elicited a P_D_ between 260 and 370 ms (z_crit_ = 1.43; P_Dmax_ = 5.43 at 320 ms; all *P* < .02; see Figure 3A). The P_D_ amplitude was not correlated with participants’ performance either, neither in a time-resolved manner (*r*_max_ = .34 at 324 ms; all *P* > .19; see Figure 3B), nor for averaged P_D_ response (*r* = .37; *P* = .14; see Figure 3B). We found the amplitude of the N_T_ twice as large as the P_D_ (*M*_NT_ = 10.80; *M*_PD_ = 4.22; *t*_16_ = 3.55; *p* = .003; see Figuer 3A). Furthermore, the N_T_ response (∼ 246 ms; *SD* = 27 ms) peaked significantly earlier compared to the P_D_ response (∼ 328 ms; *SD* = 23 ms; *t*_16_ = 10.44; *P* < .0001).

**Figure 3.**
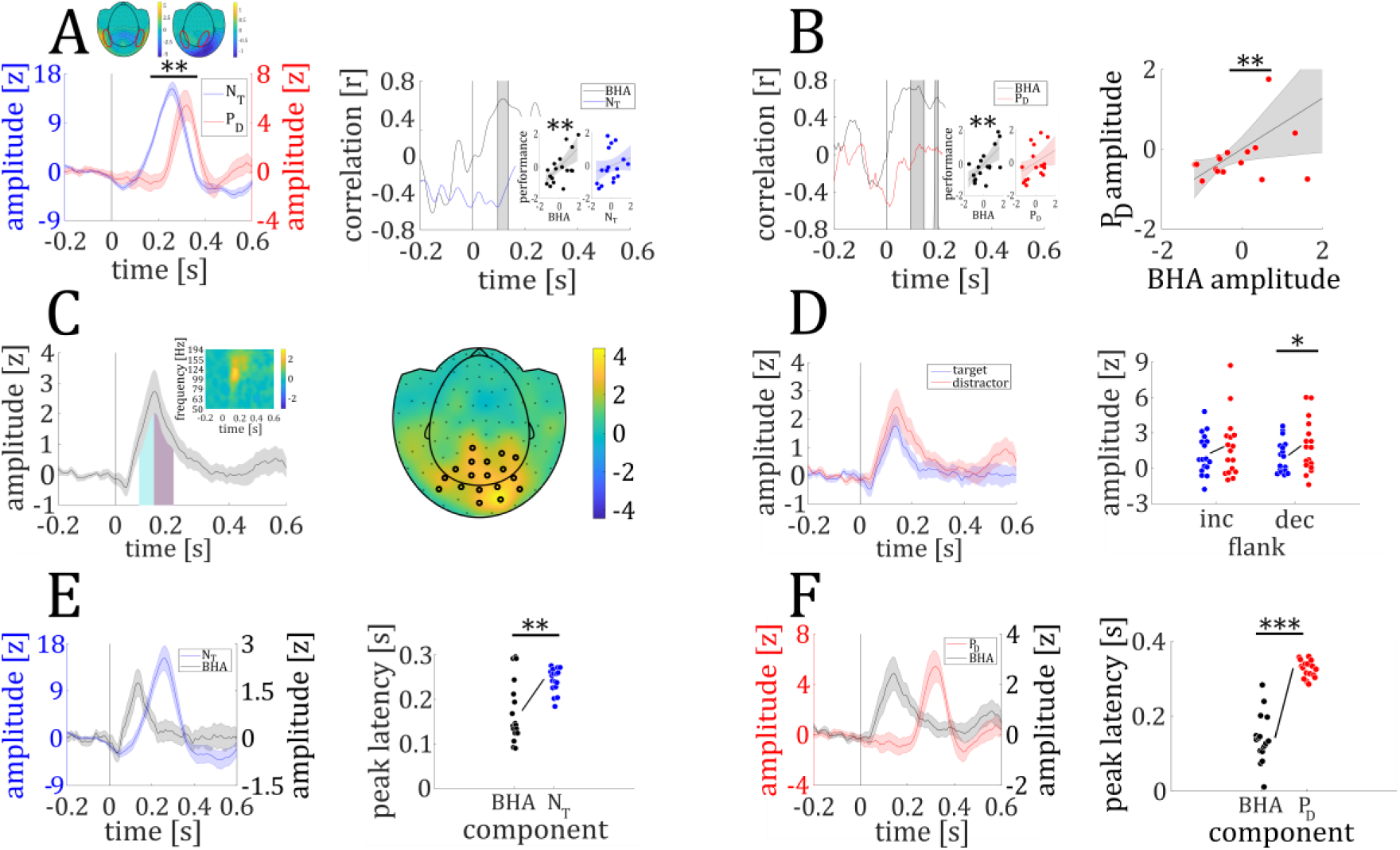
MEG results. ***A***: Time course of the NT (blue) and PD (red) responses to lateral targets and distractors, respectively (left panel). Mean amplitude of the NT response was higher compared to the PD. Upper left (NT) and right (PD) inset show the topographical distribution of MEG activity to lateral targets and distractors, respectively, with corresponding efflux / influx zones. Correlation coefficient at each time point between participants’ BHA response to lateral targets (black) and their performance and between participants’ NT (blue) and their performance (right panel). Insets show correlation coefficient between participants’ performance and their mean BHA and NT, respectively. ***B***: Correlation coefficient at each time point between participants’ BHA response to lateral distractors (black) and their performance and between participants’ PD (red) and their performance (left panel). Insets show correlation coefficient between participants’ performance and their mean BHA and PD, respectively. Maximal correlation between PD (at 344 ms) and peak BHA response (at 122 ms) (right panel). ***C***: Time course of overall BHA response, showing time intervals of increasing (cyan) and decreasing (purple) flank (left panel) and its time-frequency representation (small inset). Topographical distribution of overall BHA response with sensors showing a significant response being highlighted (right panel). ***D***: Time course of BHA response to lateral targets (blue) and distractors (red) (left panel). Individual mean BHA amplitude for lateral targets and distractors for the increasing and decreasing BHA flank (right panel). BHA amplitude of the decreasing flank was higher for lateral distractors compared to targets. ***E***: Time course for BHA to lateral targets (black) and NT (blue) (left panel). Individual time points of peak BHA and NT response (right panel). ***F***: Time course for BHA to lateral distractors and PD (third panel). Individual time points of peak BHA and PD response (right panel). Colored circles represent single data points. Shaded colored lines represent the standard error of the means (SEM). * = P < .05. ** = P < .01. *** = P < .001.

### III – Broad Band High Frequency Activity

A total of 18 occipital magnetometers showed a significant BHA response to lateral stimuli between 84 and 204 ms (z_crit_ = 0.85; BHA_max_ = 2.73 at 136 ms; all *P* values < .05; see Figure 3C).

### IV – BHA Response to Targets and Distractors

Next, we tested whether lateral targets and distractors elicited a BHA modulation. We found a significant BHA response to lateral targets between 96 ms and 180 ms (z_crit_ = 0.57; BHA_max_ = 1.76 at 134 ms; all *P* < .05; see Figure 3D) and a significant BHA response to lateral distractors between 92 and 206 ms (z_crit_ = 0.99; BHA_max_ = 2.43 at 142 ms; all *P* < .04; see Figure 3D). The increasing flank (84 ms – 134 ms, see Figure 3C) of the BHA responses did not show differences between lateral targets and distractors (*M*_Target_ = 1.27; *M*_Distractor_ = 1.84; *t*_16_ = 1.73; *P* = .10). However, BHA amplitude of the decreasing flank (134 ms – 204 ms) was higher for lateral distractors compared to lateral targets (*M*_Target_ = 1.10; *M*_Distractor_ = 1.92; *t*_16_ = 2.31; *P* = .035; see Figure 3D).

### V – Comparison between BHA Response and N_T_ / P_D_ Response

In a following step, we compared the response characteristics of the low-frequency target enhancement and distractor suppression components with the BHA’s response characteristics. *BHA response to lateralized targets vs. N_T_.* In contrast to the low-frequency N_T_ amplitude, the BHA amplitude to lateral targets was correlated to individual performance both for the time resolved analysis (*r*_max_ = .67 at 118 ms; all *P* < .0125; see Figure 3A) and the averaged BHA to targets (*r* = .65; *P* = .005; see Figure 3A). The BHA (∼ 174 ms; *SD* = 70 ms) peaked earlier than the N_T_ (∼ 246 ms; *SD* = 27 ms; *t*_16_ = 3.72; *P* = .002; see Figure 3E). Neither mean, nor peak BHA response were correlated to mean (*r* = -.08; *P* = .76) and peak (*r*_max_ = .33 at 168 ms; *P* = .19) N_T_ response, respectively.

#### BHA response to lateralized distractors vs. P_D_

The BHA to lateral distractors (at ∼ 142 ms; *SD* = 64 ms) peaked earlier than the P_D_ (∼ 328 ms; *SD* = 23 ms; *t*_16_ = 7.51; *P* < .0001; see Figure 3F). Both mean and peak BHA response were correlated to mean (*r* = .57; *P* = .017) and peak (*r*_max_ = .64 at 344 ms; *P* = .006; see Figure 3B) P_D_ response, respectively. The BHA amplitude to lateralized distractors, but not the P_D_, was correlated to performance (time-resolved analysis: *r*_max_ = .74 at 122 ms; all *P* < .0125; average BHA amplitude: *r* = .70; *P* = .002; see Figure 3B).

### VI – Lateralized BHA response

Seven stimulus-responsive channels were located in the left hemisphere and nine in the right hemisphere (see *Results III – Broadband High Frequency Activity*; see Figure 4A). When we compared mean contralateral and ipsilateral grand average BHA responses in the time range of the decreasing flank, we found a higher BHA response contralateral to the presented stimulus compared to ipsilateral (*M*_contra_ = 2.47; *M*_ipsi_ = 1.64; *t*_16_ = 2.22; *P* = .04; see Figure 4B). BHA response to lateral targets did not significantly differ between contralateral and ipsilateral hemisphere (*t*_16_ = 1.24; *P* = .23; see Figure 4C), neither did BHA response to lateral distractors (*t*_16_ = 0.61; *P* = .55; see Figure 4D).

**Figure 4.**
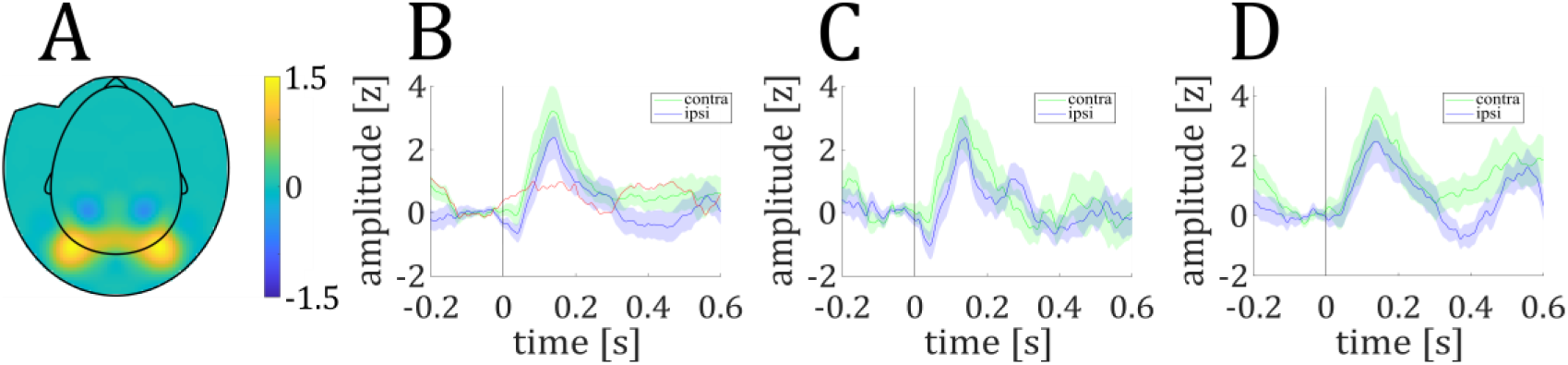
Lateralized BHA Response. ***A***: Topographical distribution of overall BHA response (contralateral minus ipsilateral). ***B***: Contralateral and ipsilateral BHA response to lateral stimuli (target and distractor). Red line shows the difference wave (contralateral minus ipsilateral). ***C***: Contralateral and ipsilateral BHA response to lateral targets. ***D***: Contralateral and ipsilateral BHA response to lateral distractors. Shaded colored lines represent the standard error of the means (SEM).

### VII Behavioral Consequences of Implicit Learning

#### Discrimination performance

Performance did not differ between learning and non-learning conditions, neither for all lateral distractor trials (*M*_learning_ = 80.5 %; *M_non-learning_* = 80.2 %; *t_29_* = 0.19; *P* = .85), nor for trials where the distractor was presented at the position with the highest occurrence probability (*M*_learning_ = 80.4 %; *M_non-learning_* = 79.6 %; *t_29_* = 0.53; *P* = .60; see Figure 5B).

**Figure 5.**
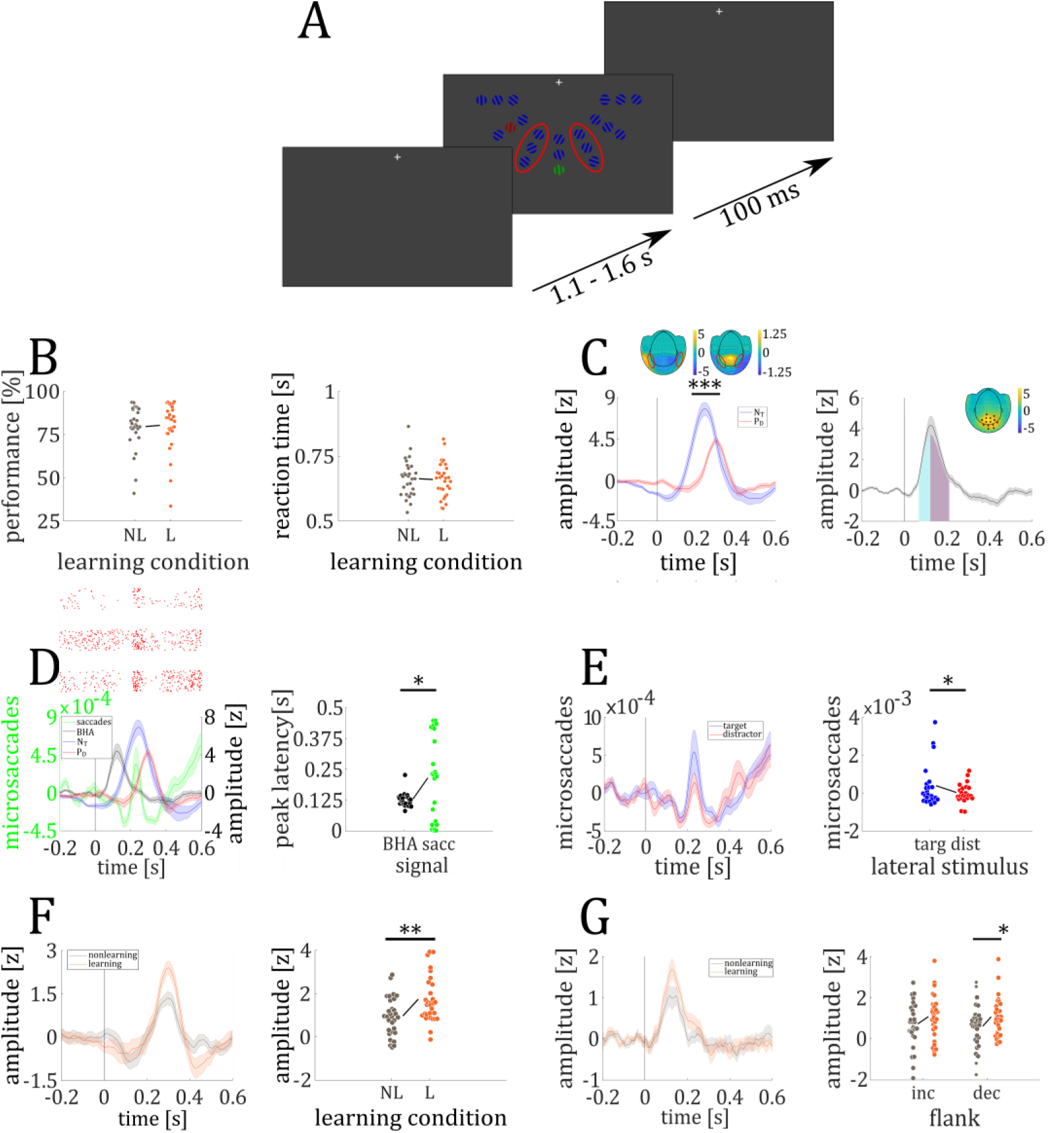
Implicit Learning Experiment. ***A***: Depiction of the paradigm. Red ellipses (not visible in experiment) mark distractor positions closest to the midline used for statistical learning. ***B***: Performance (left panel) and reaction times (right panel) did not differ between learning and non-learning trials. ***C***: Time course of the NT (blue) and PD (red) grand average responses to lateral targets and distractors, respectively (left panel). The NT response was higher compared to the PD response. Upper left (NT) and right (PD) inset show the topographical distribution of MEG activity to lateral targets and distractors, respectively, with corresponding efflux / influx zones. Time course of overall BHA response, showing time intervals of increasing (cyan) and decreasing (purple) flank (right panel). Topographical distribution of overall BHA response with sensors showing a significant response being highlighted (small inset). ***D***: Time course of microsaccadic eye movements (< 0.5° va), BHA, NT and PD response (left panel). Red dots mark single microsaccadic events of three example participants (upper inset, 3 rows). Individual time points of peak BHA and microsaccade response (right panel). ***E***: Time course of microsaccades to lateral targets and distractors (left panel). Individual mean microsaccade rate to lateral targets and distractors (right panel). ***F***: Time course of PD response in learning and non-learning condition (left panel). Individual mean PD amplitude for learning and non-learning trials, for distractors close to the midline. Mean PD response was higher in learning compared to non-learning trials (right panel). ***G***: Time course of BHA response in learning and non-learning condition (left panel). Individual mean BHA amplitude for learning and non-learning trials for the increasing and decreasing BHA flank (right panel). BHA amplitude of the decreasing flank was higher for learning trials compared to non-learning trials. Colored circles represent single data points. Shaded colored lines represent the standard error of the means (SEM). * = *P* < .05. ** = *P* < .01. *** = *P* < .0001.

#### Reaction times

Reaction times did not differ between learning and non-learning conditions, neither for all lateral distractor trials (*M*_learning_ = 659 ms; *M_non-learning_* = 661 ms; *t_29_* = 0.55; *P* = .59), nor for trials where the distractor was presented at the position with the highest occurrence probability (*M*_learning_ = 660 ms; *M_non-learning_* = 663 ms; *t_29_* = 0.69; *P* = .50; see Figure 5B).

### VIII – Stimulus Response

In experiment 2, lateral targets elicited a N_T_ response between 154 and 328 ms (z_crit_ = 1.17; N_Tmax_ = 7.74 at 244 ms; all *P* < .05; see Figure 5C), while lateral distractors elicited a P_D_ response between 234 and 362 ms (z_crit_ = 0.59; P_Dmax_ = 4.29 at 298 ms; all *P* < .003; see Figure 5C). Furthermore, the N_T_ showed a stronger modulation than the P_D_ response (*M*_NT_ = 5.44; *M*_PD_ = 2.97; *t*_29_ = 4.63; *P* = .00007; see Figure 5C).

A total of 14 occipital magnetometers showed a significant BHA response to lateral stimuli between 68 and 210 ms (z_crit_ = 0.60; BHA_max_ = 4.22 at 122 ms; all *P* values < .05; see Figure 5C).

### IX – Temporal Evolution of Microsaccades

We compared microsaccade, BHA and low-frequency N_T_ and P_D_ peak times. We found no difference in peak times between microsaccades and N_T_ (microsaccades_peak_ = 215 ms; NT_peak_ = 251 ms; *t*_22_ = 0.98; *P* = .34), we found that the P_D_ peaked later than microsaccades (PD_peak_ = 304 ms; *t*_22_ = 2.39; *P* = .026). In contrast we found that BHA precedes microsaccades (microsaccades_peak_ = 215 ms; BHA_peak_ = 131 ms; *t*_22_ = 2.40; *P* = .025; see Figure 5D).

We found more microsaccades to lateral targets compared to distractors (*M*_Target_ = 0.0004; *M*_Distractor_ = 0.00003; *t*_22_ = 2.20; *P* = .038; see Figure 5E) indicating that differences in amplitude modulation of N_T_ (*M* = 5.44) and P_D_ (*M* = 2.97) might at least in part be explained by differences in microsaccade rate.

### X – Amplitude Modulation with Experimental Condition

When we compared mean P_D_ responses between learning and non-learning trials for distractors located close to the midline (marked positions in Figure 5A), we found a higher P_D_ response in learning trials compared to non-learning trials (*M*_learning_ = 1.74; *M*_non-learning_ = 0.97; *t*_29_ = 2.88; *P* = .007; see Figure 5F). The target selection, as indexed by the N_T_, showed no difference between learning and non-learning trials (*M*_learning_ = 2.17; *M*_non-learning_ = 2.09; *t*_29_ = 0.30; *P* = .77), which was expected since the statistical probability of the target locations was identical for learning and non-learning blocks.

While the increasing flank of the BHA (68 – 122 ms) showed no significant difference between learning and non-learning trials (*M*_learning_ = 1.08; *M*_non-learning_ = 0.73; *t*_29_ = 1.44; *P* = .16), we found the decreasing flank of the BHA (122 – 210 ms) in learning trials to be increased in amplitude compared to non-learning trials (*M*_learning_ = 1.07; *M*_non-learning_ = 0.61; *t*_29_ = 2.06; *P* = .0485; see Figure 5G).

## Discussion

We examined whether broadband high-frequency activity (BHA) is an early marker of spatial selective attention and distractor suppression. Specifically, we compared the BHA response in the MEG to low-frequency N_T_ and P_D_ components. We found that the BHA recorded at MEG sensors covering the occipital cortex distinguished between lateral targets and distractors with a higher amplitude to lateral distractors. The BHA preceded the low-frequency components by more than 100 ms. Most importantly, the early BHA modulation was correlated to individual performance and P_D_ (but not N_T_) response. In a second experiment, we examined how the P_D_ and BHA responses are affected by implicit learning and compared the time course of microsaccades to BHA, N_T_ and P_D_. Both, the P_D_ and BHA to lateral distractors, increased in amplitude during implicit learning indicative of a trainable distractor suppression mechanism. Microsaccades peaked later than the BHA, but in the time range of the low-frequency components. These results indicate that the early visual BHA reflects the initial instance of effective distractor suppression.

In general, we found better target discrimination with higher target orientation angle. This is in line with previous literature showing a performance increase with increasing target angle (Wienke et al., 2021). Participants discriminated central targets better than lateral targets. This location-based effect is in line with a previous study investigating target and distractor processing using a visual search task (Hilimire et al., 2011). Participants were presented with a search paradigm consisting of 16 letters comprising targets (colored horizontally/upright “T”), distractors (colored horizontally/upright “L”) and non-targets (gray horizontally “T”). This paradigm could not identify a feature-based distractor effect because target and distractor similarity was not systematically varied. However, consistent with our study the authors found faster reaction times and lower error rates to central compared to lateral targets. Our results therefore show a clear location-dependent distractor effect.

In addition, in our study the angular difference between targets and distractors showed a feature-based distractor effect. Performance decreased with greater orientation difference between target and distractor angle. Previous studies did not address this discrimination difficulty, resulting in a lack of comparable findings (Hickey et al., 2009; Hilimire et al., 2012).

Both the N_T_ and P_D_ were elicited by lateral targets and distractors, respectively. We observed an N_T_ onset around 200 ms and a peak response around 300 ms after stimulus presentation, which is in line with previous findings (Donohue et al., 2018; Drisdelle & Eimer, 2021; Hickey et al., 2009). Critically, the P_D_ in our experiments peaked later than the N_T_ (∼80ms), but showed the same topography as found in Donohue et al. (2018). The P_D_ resembled an inverted N2pc field with an efflux component over central occipital areas and an influx component at lateral occipital areas. Despite the differences in stimuli in previous studies, we found a similar relationship between N_T_ and P_D_ magnitude. Donohue et al. (2018) used a symmetric search array with T letters around a fixation cross, restricting the color-defined target or distractor to lateral positions. In line with Donohue et al. (2018), we found that the neural response to targets (N_T_) is stronger than the response to distractors (P_D_). Given that distractor suppression is an essential process, one might expect its neural correlate to be comparable with the target selection response. Previous work also suggests that the microsaccade rate modulates the EEG/MEG amplitude (Yuval-Greenberg et al., 2008). Lateral targets require a spatial shift of attention for successful discrimination, which explains the higher microsaccades rate (Rucci et al., 2007). The spikes elicited by the microsaccadic movements could then add to the MEG amplitude (Liu et al., 2023; Lowet et al., 2018). Microsaccades in our study peaked after 200 ms (Yuval-Greenberg et al, 2008). In this time range microsaccades overlap with low-frequency components and explain their field distribution (Carl et al., 2012; Liu et al., 2023). Given that microsaccade rate to lateral targets is higher than to lateral distractors, the higher amplitude of the N_T_ might partially be explained by the presence of microsaccades. These results further indicate that the low-frequency responses might be influenced by microsaccadic eye movements. The late and reduced P_D_ in addition to fewer microsaccades compared to target trials indicate that the suppression process has already been initiated earlier.

The BHA showed a stronger modulation to lateral distractors compared to targets. The peak BHA predicted the strength of individual P_D_ (but not N_T_) responses, which is in line with the cortical “selection for rejection” mechanism (Bartsch et al., 2021; Donohue et al., 2018). The authors proposed the N1pc as an early selection for rejection marker. The N1pc peaked around the decreasing flank of the BHA in our study. However, in contrast to the BHA, the N1pc did not show differences in amplitude to lateral targets versus distractors. Hence, the N1pc can be seen as an a general “attend-to-me” signal to salient items (Sawaki & Luck, 2010). Since the BHA was only correlated to the P_D_, but not the N_T_ our results suggest that the BHA represents a more selective “selection for rejection” mechanism and thus serves as an early marker of distractor suppression.

BHA has been the focus of studies using human intracranial recordings (Miller et al., 2014) but is only recently described in non-invasive recordings (Wienke et al., 2021). Intracranial studies demonstrated how BHA varies in different visual areas (Bartoli et al., 2019; Gerber et al., 2017; Golan et al., 2017; Vishne et al., 2023). Our observed BHA response with a fast onset prior to 100 ms, a peak prior to 200 ms and a slowly decreasing flank closely resemble the BHA observed in these studies (Bartoli et al., 2019; Gerber et al., 2017; Golan et al., 2017; Vishne et al., 2023), especially the BHA response in V1 electrodes (Bartoli et al., 2019; Golan et al., 2017).

Previous intracranial recordings in epilepsy patients found the BHA to be increased during attentional selection of auditory stimuli (Ray et al., 2008). In the visual domain, previous studies investigating attentional modulation of activity primarily focused on broad gamma-bands between 30 and 130 Hz with mixed results (Davidesco et al., 2013; Tallon-Baudry et al., 2005). Tallon-Baudry et al. (2005) found an increase in baseline gamma-range activity (30 – 130 Hz) and a decrease during stimulus presentation in lateral occipital cortex, but an increase over fusiform gyrus during attentive states. In contrast, Davidesco et al. (2013) presented participants with small and large objects and cued them to attend one of the stimuli. They found activity in the gamma-range (30 – 90 Hz) in visual cortex to be increased during attentive states, while previous studies using a narrower gamma-range (30 – 50 Hz) failed to show such modulations (Chalk et al., 2010). Furthermore, Davidesco et al. (2013) found attentional modulation of gamma activity to occur earlier in early visual cortical areas compared to higher visual areas. High-frequency modulation (> 80 Hz) during selective attention was also found in intracranial recordings (Szczepanski et al., 2014). Participants were instructed to covertly attend either to the left or right visual field and respond to a target. Szczepanski et al. (2014) found an increase in BHA during attentive states over visual areas. Most interestingly, ∼20 % of the recorded electrodes exhibited stronger BHA responses contralateral to the attended visual field, which is in line with our results of an increased BHA in MEG sensors contralateral to lateral stimuli.

A central finding of our study is that BHA is modulated by implicit learning, a relationship not yet demonstrated. In general, spatial proximity between targets and distractors increases interference (Hickey & Theeuwes, 2011; Matho t et al., 2010; Mounts, 2000). Therefore, we increased the occurrence probability of distractors at positions close to the central targets to test whether the BHA/P_D_ responses could be modified by statistical learning. Research has shown that while individuals can detect and learn statistical regularities in their environment, this learning does not always translate into enhanced performance (Conway & Christiansen, 2006; Fiser & Aslin, 2001; Siegelman et al., 2018). In line with these results, participants in our study showed no performance improvement. We attribute this to the target orientation angles being too small to discriminate despite successful distractor suppression. Our results indicate that the BHA to lateral distractors and the later P_D_ are enhanced by implicit statistical learning. This pattern cannot be explained by a repetition suppression effect, where neural responses decrease with repetition, as the BHA was actually increased (Eckert et al., 2022). These findings suggest that BHA is a selection-for-rejection mechanism that can be reinforced by learning.

## Conclusion

Our findings extend prior work on the human BHA and spatial attention. The MEG-BHA in our study showed response characteristics strongly similar to that observed in non-human primate and human intracranial recordings. The BHA distinguished between lateralized targets and distractors and preceded the low-frequency components by more than 100 ms. Furthermore, the BHA predicted participants’ performance and P_D_ response. These results indicate that the BHA is best suited to test the temporal evolution of stimulus discrimination as a key marker of successful subsequent distractor suppression.

## Author contributions

M.B. and S.D. conceived and designed the experiment. P.S. collected the MEG data. P.S., C.R., and S.D. analyzed the data. P.S., C.R., M.B. and S.D. interpreted the data. P.S., C.R., M.B. and S.D. wrote the manuscript.

## Supporting information

Post-hoc t-tests comparing performance for the different target angles.

## Acknowledgements

This study was partially supported by the German Research Foundation - DFG grant DFG SFB-1436, TPA03. The authors thank Lena Vogelgesang for coding the experiment and recording a part of the data.

