## Supplementary material for "Broadband high frequency Activity initializes Distractor Suppression": Post-hoc t-tests comparing performance for the different target angles.

| **Target angles compared** | ***t* value** | ***P*** |
| --- | --- | --- |
| 1.5° vs. 3° | 4.43 | **.0004** |
| 1.5° vs. 4.5° | 5.66 | **.00004** |
| 1.5° vs. 6° | 8.54 | **.0000002** |
| 1.5° vs. 7.5° | 13.59 | **.0000000003** |
| 1.5° vs. 9° | 14.35 | **. 0000000001** |
| 1.5° vs. 10.5° | 16.94 | **. 00000000001** |
| 1.5° vs. 12° | 16.49 | **. 00000000002** |
| 1.5° vs. 13.5° | 18.32 | **.000000000004** |
| 1.5° vs. 15° | 15.82 | **.00000000003** |
| 3° vs. 4.5° | 4.30 | **.0006** |
| 3° vs. 6° | 7.78 | **.0000008** |
| 3° vs. 7.5° | 14.34 | **.0000000001** |
| 3° vs. 9° | 12.34 | **.000000001** |
| 3° vs. 10.5° | 15.99 | **.00000000003** |
| 3° vs. 12° | 14.57 | **.0000000001** |
| 3° vs. 13.5° | 17.01 | **.00000000001** |
| 3° vs. 15° | 19.06 | **.000000000002** |
| 4.5° vs. 6° | 1.51 | .15 |
| 4.5° vs. 7.5° | 4.88 | **.0002** |
| 4.5° vs. 9° | 4.89 | **.0002** |
| 4.5° vs. 10.5° | 5.98 | **.00002** |
| 4.5° vs. 12° | 8.01 | **.0000006** |
| 4.5° vs. 13.5° | 7.75 | **.0000008** |
| 4.5° vs. 15° | 9.46 | **.00000006** |
| 6° vs. 7.5° | 4.55 | **.0003** |
| 6° vs. 9° | 4.63 | **.0003** |
| 6° vs. 10.5° | 6.88 | **.000004** |
| 6° vs. 12° | 8.75 | **.0000002** |
| 6° vs. 13.5° | 9.12 | **.00000009** |
| 6° vs. 15° | 12.50 | **.000000001** |
| 7.5° vs. 9° | 2.15 | .05 |
| 7.5° vs. 10.5° | 4.92 | **.0002** |
| 7.5° vs. 12° | 5.67 | **.00004** |
| 7.5° vs. 13.5° | 4.91 | **.0002** |
| 7.5° vs. 15° | 6.30 | **.00001** |
| 9° vs. 10.5° | 2.35 | .03 |
| 9° vs. 12° | 3.28 | .005 |
| 9° vs. 13.5° | 2.47 | .03 |
| 9° vs. 15° | 3.08 | .007 |
| 10.5° vs. 12° | 2.12 | .05 |
| 10.5° vs. 13.5° | 0.71 | .49 |
| 10.5° vs. 15° | 1.72 | .11 |
| 12° vs. 13.5° | 1.34 | .20 |
| 12° vs. 15° | 0.18 | .86 |
| 13.5° vs. 15° | 0.92 | .37 |

***Table 1***. Post-hoc t-tests comparing performance for the different target angles.
